# Distinct Mechanisms of Surround Modulation for Dynamic Natural Scenes in Motion-and Scene-Selective Cortex

**DOI:** 10.1101/2025.11.06.686956

**Authors:** Merve Kiniklioglu, David Pitcher, Daniel Kaiser

## Abstract

Surround modulation refers to changes in neural responses to a central stimulus induced by its surrounding context, which can manifest as either suppression or facilitation. Although this phenomenon is well established for simple stimuli, its underlying neural mechanism during natural scene perception remains unclear. Using fMRI, we examined how motion congruence and categorical similarity between central and surrounding scenes shape surround modulation across the visual hierarchy. Central and surrounding scenes systematically varied in categorical similarity (identical exemplar, different exemplar of the same basic-level category, different basic-level category, or different superordinate category) and in motion direction (same or opposite direction). Neural responses were measured in primary visual cortex (V1), motion-selective cortex (hMT+), and scene-selective occipital and parahippocampal place areas (OPA and PPA). hMT+ showed robust motion-dependent surround suppression, which was stronger for same-direction motion. In contrast, V1 showed surround facilitation across all conditions, which was reduced when center and surround were identical and moved in the same direction, consistent with sensitivity to physical similarity. OPA and PPA primarily exhibited facilitation. Multivariate decoding between center-only and center-surround conditions complemented these findings, revealing motion-dependent surround modulation in hMT+, and category-dependent surround modulation in OPA and PPA. Across the visual hierarchy, surrounding scene context thus systematically attenuates redundant input or enhances informative differences: from low-level facilitation in V1 to motion-dependent suppression in hMT+ and category-dependent modulation in scene-selective regions.

## Introduction

Sensitivity to visual stimuli is strongly influenced by the surrounding context. A well-known example is surround modulation, in which the response of a neuron to a stimulus presented in the receptive field (RF) center is modulated by stimuli presented in the surround. This modulation can manifest as suppression (*surround suppression*) or facilitation (*surround facilitation*) (Cavanaugh et al., 2002a, 2002b; DeAngelis et al., 1994; Ichida et al., 2007; Petrov et al., 2005; Pihlaja et al., 2008; Shushruth et al., 2012; Walker et al., 1999; Williams et al., 2003; Xing & Heeger, 2001; Zenger-Landolt & Heeger, 2003). Such modulatory effects are generally attributed to antagonistic center–surround interactions within the visual cortex, particularly in motion-selective regions such as the middle temporal area (MT) (e.g., MT hypothesis; see Er et al., 2020; Kiniklioglu & Boyaci, 2025; Pack et al., 2005; Schallmo et al., 2018; Tadin et al., 2003, 2011; Turkozer et al., 2016).

These center–surround mechanisms are observed throughout the visual hierarchy. Surround modulation has been classically demonstrated in retinotopic areas of the early visual cortex (V1-V4) and in motion-selective regions such as MT and MST, suggesting that it reflects a canonical computation of contextual modulation within the visual system (Allman et al., 1985; Angelucci et al., 2017; Hallum & Movshon, 2014; Nurminen & Angelucci, 2014; Sundberg et al., 2009). More recently, surround modulation has also been observed in higher-level regions of the ventral visual stream, such as the lateral occipital complex (LOC), highlighting its prevalence across the visual hierarchy (Montoya et al., 2025).

The strength and form of surround modulation depend on stimulus properties. Both neural (Kiniklioglu & Boyaci, 2025; Schallmo et al., 2018; Shushruth et al., 2013; Turkozer et al., 2016; Zenger-Landolt & Heeger, 2003) and behavioral studies (Er et al., 2020; Kiniklioglu & Boyaci, 2022; Schallmo et al., 2018; Tadin et al., 2011; Turkozer et al., 2016) show that low-level features such as size, contrast, and motion direction systematically influence the strength of surround suppression. Typically, feature similarity between center and surround increases suppression, whereas dissimilarity reduces it and can even reverse the effect into facilitation. For example, neurons responding to a central grating are strongly suppressed by iso-oriented surrounds but can be facilitated when the surround is orthogonal (Cavanaugh et al., 2002b; Schallmo et al., 2016; Serrano-Pedraza et al., 2012). A comparable pattern is observed for motion, where suppression is typically stronger when the center and surround drift in the same direction than when they move in opposite directions (Allman et al., 1985; Born & Tootell, 1992; Cavanaugh et al., 2002b; Kiniklioglu & Boyaci, 2025; Lamme, 1995; Paffen, van der Smagt, et al., 2005).

Although these effects are well established with simple stimuli such as static or drifting gratings, it remains unclear whether surround modulation also occurs in the context of complex, dynamic scenes. Our recent behavioral work demonstrated that surround modulation is indeed observed with dynamic natural scenes, with suppression increasing as the categorical similarity between the center and surround scenes increases (Kiniklioglu & Kaiser, 2025). These results align with scene categorization studies showing that categorically congruent surrounds facilitates recognition, whereas incongruent or physically mismatched surrounds impair it (Faurite et al., 2024; Peyrin et al., 2021). Consistent with low-level studies (Kiniklioglu & Boyaci, 2022; Paffen, Alais, & Verstraten, 2005; Paffen, van der Smagt, et al., 2005), we also found that suppression decreased when the center and surround moved in opposite directions. Together, this evidence suggests that surround modulation in natural vision can be shaped by both low-level visual features, such as motion direction, and higher-level contextual information, such as categorical similarity.

While our behavioral results suggest a suppressive effect that scales with the dissimilarity between center and surround content, neurophysiological work has disagreed on whether suppression or facilitation prevails. In the macaque primary visual cortex (V1), suppression was stronger when the center and surround were contained homogeneous natural images (Coen-Cagli et al., 2015), whereas in the cat visual cortex, surrounds that perceptually completed a central pattern elicited facilitation (Onat et al., 2013). However, no human studies have investigated the neural mechanisms underlying the suppressive effects observed in behavioral studies with dynamic natural scenes.

To address this gap, we used functional magnetic resonance imaging (fMRI) to investigate the neural correlates of surround modulation in dynamic natural scenes. We examined responses across regions of interest spanning multiple levels of the visual hierarchy, including primary visual cortex (V1), motion-selective cortex (hMT+), and the scene-selective occipital place area (OPA) and parahippocampal place area (PPA). Participants viewed central scenes presented together with surrounding scenes that varied in categorical similarity across four levels: identical exemplar, different exemplar from the same basic-level category, different basic-level categories within the same superordinate category, and different superordinate categories. This manipulation enabled us to test how categorical similarity shapes neural signatures of surround modulation across the visual hierarchy. We also varied motion congruence, comparing conditions in which the center and surround drifted in the same versus opposite directions, to assess whether motion-related effects observed with simple stimuli generalize to naturalistic stimuli. By independently manipulating categorical similarity and motion congruence, we could test how both high- and low-level contextual factors influence surround modulation in dynamic natural scenes.

## Methods

### Participants

Twenty healthy volunteers (11 female, mean age = 26.7 years) participated in the study. One participant was excluded due to excessive head motion during scanning (see Preprocessing for motion-quality criteria), leaving a final sample of nineteen participants. The sample size (N = 19) is comparable to that used in previous studies investigating low-level surround modulation (Er et al., 2020; Kiniklioglu & Boyaci, 2025; Schallmo et al., 2016). All reported normal or corrected-to-normal vision. Written informed consent was obtained before participation, and participants received monetary compensation. The study was approved by the Ethics Committee of Justus Liebig University Giessen. All experimental protocols were in accordance with the Declaration of Helsinki.

### Stimuli

Stimuli were panoramic videos created by moving static scene images behind a circular occluder (908 × 699 pixels; 5 pixels per frame). The central image was presented through a 1.9°-diameter aperture, surrounded by an annulus ranging from 2.5° to 10.4°. To separate the center and surround, the area between them (1.9°-2.5°) remained unstimulated. A cosine envelope was applied at the aperture boundaries to minimize sharp transitions at the edges.

Scene images were selected from two superordinate categories (indoor: restaurants, museums; outdoor: parks, residential areas), each featuring two basic-level categories, with two exemplars per basic-level category. The center scenes were always presented with a surrounding scene, except in center-only trials where the central stimulus appeared alone (see below). The resulting center–surround stimuli depicted natural scenes that varied in their categorical relationship between center and surround, with four conditions: identical exemplar condition, where the center and surround images were identical; different exemplar condition, where the center and surround images belonged to the same basic-level category (e.g., museums) but were not identical; different basic-level category condition, where the center and surround images belonged to the same superordinate category (e.g., indoor) but were from different basic-level categories; and different superordinate category condition, where the center and surround images belonged to different superordinate categories (see Figure 1). Motion congruency was manipulated with two conditions: in half of the trials, the center and surround scenes moved in the same direction, and in the other half, they moved in opposite directions.

**Figure 1:**
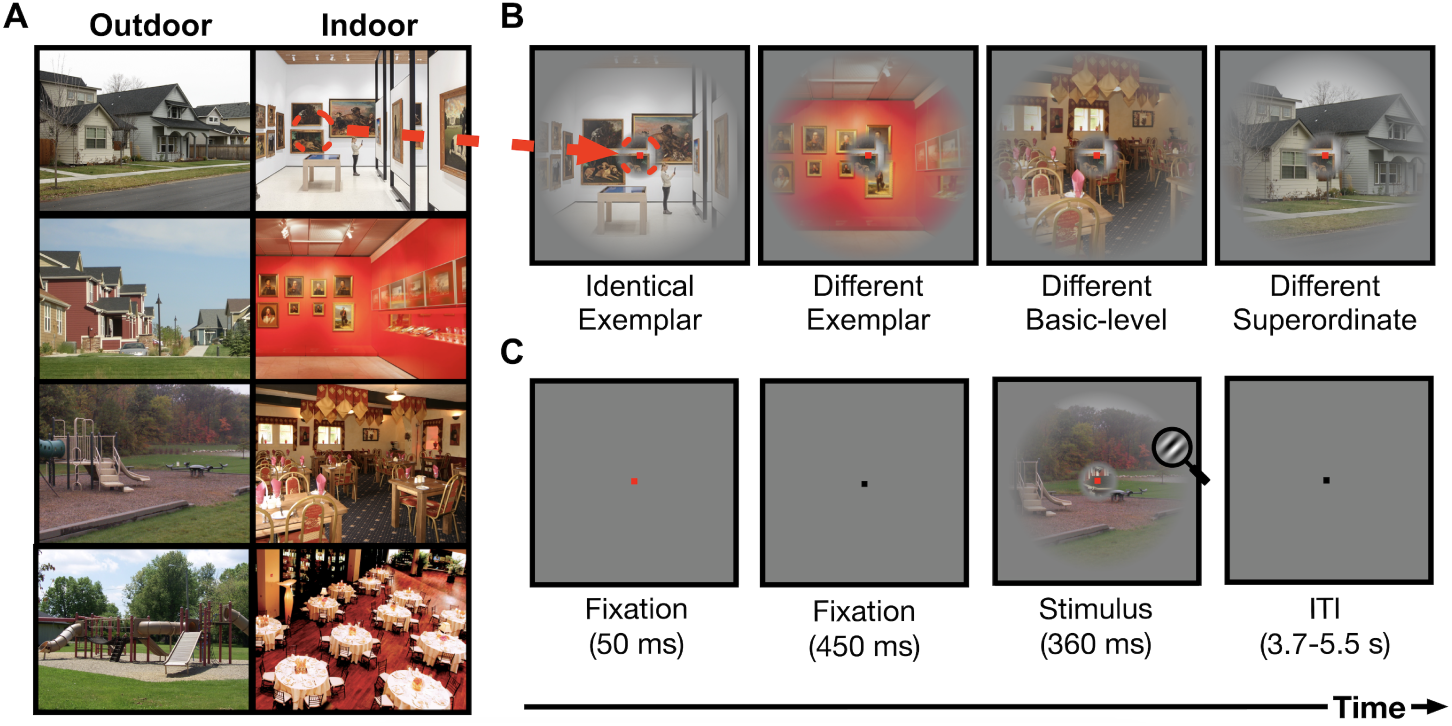
Stimuli and Paradigm. A) The natural scene images used to create the stimuli were drawn from two superordinate categories (indoor and outdoor) and two basic-level categories within each (restaurants, museums, parks, and residential areas), with two exemplars per basic-level category. B) Example stimuli for each category condition, shown from left to right: identical exemplar, different exemplar, different basic-level category, different superordinate category. C) Experimental design. Each trial began with a fixation point (500 ms), which briefly turned red to signal the upcoming stimulus, followed by presentation of a scene video (360 ms). Intertrial intervals varied randomly between 3.7 and 5.6 s. On 10% of trials, a high-contrast Gabor patch appeared at a random location within either the center or surround region, and participants reported its detection with a keypress.

### Experimental Paradigm

We used a mixed event-related design in which center-only and center+surround trials were presented within the same run (Figure 1). In total, nine trial types (four category conditions × two motion directions, plus the center-only condition) were presented in randomized order. Center-only trials served as a baseline for quantifying the suppressive effect of adding the surround.

Each trial began with a fixation point (500 ms), followed by a scene video (360 ms) on a mid-gray background. Participants were instructed to maintain fixation at the center of the display. Intertrial intervals varied randomly between 3.7 and 5.6 seconds, sampled from a uniform distribution. To ensure that participants attended to both center and surround, on 10% of trials a small Gabor patch (spatial frequency = 1 cycles/°, diameter = 0.7°, contrast = 98%) appeared at a random location within either the center or surround region. Participants reported its detection with a keypress, with a mean accuracy of 92.2% (SD = 7.6%). Each condition was presented eight times per run, and participants completed five runs. Each run lasted approximately 10 minutes, and short breaks were provided between runs.

### Data Acquisition

MRI data were acquired on a 3T Siemens Magnetom Prisma scanner (Siemens Healthineers, Erlangen, Germany) equipped with a 64-channel head coil at the Bender Institute of Neuroimaging (BION), Justus Liebig University Giessen. High-resolution anatomical images were collected using a T1-weighted 3D sagittal MP-RAGE sequence (voxel size = 1 mm^3^ isotropic, 176 slices). Functional images were acquired with a T2*-weighted EPI sequence (TR = 1850 ms, TE = 30 ms, voxel size = 2.2 mm^3^, 58 slices). Visual stimuli were presented on a MR-compatible LED monitor (1920 × 1080 pixels, 120 Hz) positioned at the rear of the bore and viewed via a mirror mounted on the head coil at a distance of 140 cm. Stimuli were generated and presented using MATLAB (MathWorks, Natick, MA) and the Psychophysics Toolbox (Brainard, 1997). Participant responses were recorded using an MR-compatible fiber-optic response box. Each session began with an anatomical scan, followed by four localizer runs and five experimental runs, for a total duration of approximately 90 minutes.

### Localizer Runs

*hMT+:* The hMT+ area of each participant was defined using established methods (Huk et al., 2002). Stimuli consisted of 200 randomly positioned white dots presented within a 12° aperture on a black background. In dynamic blocks, the dots moved along one of four trajectories: radial (expansion–contraction), angular (clockwise–counterclockwise rotation), horizontal (left–right), or vertical (up–down). Motion direction changed every 1.85 s to prevent adaptation. In static blocks, the dots were identical but remained fixed in position. Each block lasted 14.8 s, and dynamic and static blocks alternated eight times per run. Participants maintained central fixation and performed a color-change detection task in which they reported changes in the color of the fixation point.

*V1:* The V1 area of each participant was defined using a flickering checkerboard-patterned wedge paradigm, similar to established retinotopic mapping methods (Engel et al., 1997; Sereno et al., 1995). Because we were specifically interested in the V1/V2 boundary, we used alternating horizontal and vertical wedges instead of rotating or expanding stimuli (Greenberg et al., 2012; Slotnick & Yantis, 2003). The run consisted of 14.8-s blocks of horizontal and vertical wedges presented in alternation, repeated ten times in the run. As in the hMT+ localizer, participants maintained fixation and performed a color-change detection task.

*OPA and PPA:* Scene-selective areas were defined using 3-s movie clips from four categories: faces, scenes, objects, and scrambled objects (Küçük et al., 2024; Pitcher et al., 2011). Each category included 60 clips. Face clips featured seven children filmed against a black background in close-up, showing only their faces as they danced or interacted with toys or with adults who remained out of frame. Scene clips were recorded in 15 different locations, primarily pastoral settings, and filmed from a slowly moving car. Object clips depicted 15 distinct inanimate items (e.g., mobiles, wind-up toys, toy planes, tractors, and rolling balls). Scrambled-object clips were generated by dividing each object movie into a 15 × 15 grid and spatially shuffling the resulting segments within each frame. Participants completed eight blocks per category, each consisting of six videos randomly sampled from the full set of 60 exemplars in that category. Although the same actor, scene, or object could appear more than once, the large stimulus pool made such repetitions unlikely. As in the previous localizer runs, participants maintained fixation and reported changes in the color of the fixation point.

### hMT+ and V1 Sub-ROI Localizer Run

Using an independent localizer run, we identified the voxels corresponding to the spatial location and size of the center stimuli as sub-ROIs within hMT+ and V1. This approach is commonly used to localize responses to low-level grating stimuli (Er et al., 2020; Kiniklioglu & Boyaci, 2025; Schallmo et al., 2018), but here we adapted it to ensure that the same spatially defined sub-ROIs could be applied in the analysis of naturalistic stimuli. In the localizer, participants viewed drifting high-contrast (98%) center and surround (i.e., annulus) gratings, matched in size and location to those used in the functional runs. The run consist of 14.8-s center blocks alternating with 14.8-s surround blocks, each separated by a 14.8-s blank period, repeated six times. As in the other localizers, participants maintained fixation and reported changes in the color of the fixation point.

## Data Analysis

### Preprocessing

*Localizer Runs:* Localizer data were preprocessed and analyzed using the FMRIB Software Library (FSL) (www.fmrib.ox.ac.uk/fsl) and Freesurfer (Dale et al., 1999; Fischl et al., 1999; Woolrich et al., 2001). High-resolution anatomical images were skull-stripped with BET. Preprocessing steps for functional images included motion correction with MCFLIRT, high-pass temporal filtering (100s), and BET brain extraction. Each participant’s functional images were aligned to their own high-resolution anatomical image and registered to the standard Montreal Neurological Institute (MNI) 2-mm brain using FLIRT. The 3D cortical surface was constructed from anatomical images for each participant using FreeSurfer’s *recon-all* command for visualizing statistical maps, anatomical delineation, and identifying ROIs.

*Experimental Runs:* Experimental data were preprocessed and analyzed in SPM12 (www.fil.ion.ucl.ac.uk/spm/). Preprocessing steps included geometric distortion correction with the SPM FieldMap toolbox, motion correction and coregistration of functional volumes to each participant’s T1-weighted structural image. Structural images were segmented and normalized to MNI 2-mm standard space. After also normalizing the functional images, they were spatially smoothed with a 6 mm FWHM Gaussian kernel.

To assess data quality, head motion was quantified for each volume using framewise displacement (FD). Volumes with FD greater than 0.5 mm were flagged as motion outliers (Power et al., 2012). Runs with more than 30% flagged volumes were discarded (Parkes et al., 2018). Participants with two or more runs discarded were excluded (Ciric et al., 2017). One participant met this exclusion criterion and was removed from all analyses; all results reported below are based on the remaining *N* = 19 participants.

### ROI Construction

For all ROI constructions, a general linear model (GLM) was applied using FSL’s FMRI Expert Analysis Tool (FEAT). The predicted fMRI response in each trial was computed assuming a double-gamma hemodynamic response function (HRF). Nuisance regressors for linear motion (derived from MCFLIRT) were also included in the model. For removing temporal autocorrelations, FILM prewhitening was applied (Woolrich et al., 2001).

*hMT+:* For the hMT+ ROI, the statistical parametric maps (SPMs) of the dynamic versus static contrast were registered to Freesurfer and overlaid on the surface in the native space using the *tksurfer* command. Utilizing the MT label from FreeSurfer’s anatomical delineation for guidance, voxels at the ascending tip of the inferior temporal sulcus and responding more to dynamic compared to static dots were identified as hMT+ and used as a mask for the hMT+ sub-ROI localization.

*V1:* For the V1 ROI, SPMs of horizontal versus vertical contrast were registered to Freesurfer and overlaid on the surface in the native space using the *tksurfer* program. Utilizing the V1 label from FreeSurfer’s anatomical delineation for guidance and voxels that respond stronger to vertical than horizontal wedges, we drew the V1-V2 boundaries. The voxels that fell in or around the calcarine sulcus were identified as V1 and used as a mask for the V1 sub-ROI localization.

*hMT+ and V1 sub-ROIs:* For V1 and hMT+ sub-ROIs, we analyzed the independent localizer run data and identified the voxels that respond more strongly to the center compared to the surround grating within V1 and hMT+. The activated regions were identified as V1 and hMT+ sub-ROIs, respectively, and subsequently used as masks for the analysis of the experimental runs.

*OPA and PPA:* For the scene-selective ROIs, we identified the voxels with the greatest scene versus face and object contrast, using a voxelwise corrected threshold of *p <* 0.05, and a cluster extent threshold of 10 voxels. The occipital place area (OPA) was localized as the scene-selective cluster on the lateral surface near the transverse occipital sulcus, and the parahippocampal place area (PPA) as the scene-selective cluster on the ventral visual cortex. The identified regions were used as OPA and PPA masks. Because the center vs. surround contrast did not yield reliable voxel-level responses in OPA and PPA, we used the full ROIs rather than sub-ROIs for the analysis of the experimental runs.

### Analysis of Experimental Runs

#### Univariate Analysis

The experimental runs were analyzed in SPM12 using a general linear model (GLM). For each run, separate regressors were specified for each of the nine conditions, and six motion parameters from realignment were included as nuisance regressors. Predicted BOLD responses were modeled by convolving event onsets with a double-gamma hemodynamic response function (HRF), and statistical parametric maps (SPMs) were generated to assess the main effect of each condition across runs. To quantify fMRI responses, parameter estimates (beta weights) for each condition were extracted from the predefined ROIs (hMT+, V1, OPA, and PPA) using custom code with SPM12 functions. For these ROI-based univariate analyses we used the unsmoothed preprocessed data to preserve fine-grained voxel-level signals. The beta weights were treated as fMRI responses and used in further statistical analyses.

To quantify changes in fMRI response due to the presence of the surround, we calculated a Suppression Index (SI), defined as:

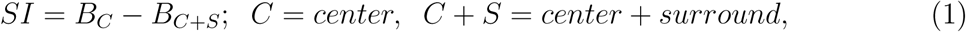

where *B.* denotes the fMRI response for the given condition. Negative SI values indicate surround facilitation, positive values indicate surround suppression, and an SI of 0 reflects no suppression or facilitation.

Surround modulation was assessed by testing SI values against zero using one-sample, two-tailed Student’s *t*-tests with correction for multiple comparisons. We then performed a two-way Analysis of Variance (ANOVA) on SI values with categorical similarity (four levels: identical exemplar, different exemplar, different basic-level category, different superordinate category) and motion-direction congruence (two levels: same-direction, opposite-direction) as factors for each sub-ROI using SPSS Version 25 (IBM Corp., Armonk, NY). Finally, post hoc paired-sample *t*-tests were used to assess how categorical similarity affects surround modulation under each motion condition.

#### Multivariate Analysis

To investigate distributed patterns of activity, we performed ROI-based decoding using CoSMoMVPA (Oosterhof et al., 2016) on condition-wise GLM parameter estimates. For each participant and ROI (hMT+, V1, OPA, and PPA), beta images from each run and condition were assembled into voxel pattern vectors. We implemented two-way classification, contrasting *center-only* with *center+surround* trials across different category and motion-direction conditions, using linear discriminant analysis (LDA). Datasets were z-scored within the training folds and class-balanced within chunks (runs). Cross-validation used a leave-one-run-out partitioning scheme, and the mean accuracy across folds was taken as the decoding score.

This two-way decoding quantifies the separability of multivoxel response patterns between center-only and center+surround conditions. Accuracy above chance indicates that the presence of a surround reliably alters the distributed activity pattern across voxels relative to center-only. Higher decoding accuracy reflects a greater surround-induced change in these patterns, either due to shifts in overall response magnitude and/or reconfiguration of voxelwise patterns, and thereby indexes a stronger influence of the surround on neural representations. We thus interpret decoding accuracy as an indicator of the magnitude of surround modulation, while it cannot indicate the direction of the modulation (i.e., suppression vs facilitation).

Group-level significance was assessed by comparing subject-level accuracies against chance level (0.5) using one-sample t-tests (one-tailed), with false discovery rate (FDR) correction applied across conditions. For all ROIs (hMT+, V1, OPA, and PPA), we performed separate decoding analyses for all combinations of scene categories and for both motion directions. For assessing the effects of categorical similarity, data were merged across motion directions to increase statistical power, and decoding accuracies were compared across the four category conditions using a one-way repeated-measures ANOVA. For assessing the effects of motion congruence, data were merged across categories, and directional sensitivity was assessed by directly comparing decoding accuracy between same- and opposite-direction conditions using paired-samples *t*-test.

## Results

### Univariate analyses reveal distinct patterns of surround suppression and facilitation across visual regions

In hMT+, we observed surround suppression for conditions in which the center and surround moved in the same direction. Figure 2A shows the Suppression Index (SI; see Methods) for the hMT+ sub-ROI across four categorical similarity and two motion-direction conditions. SI values were significantly greater than zero only in the identical-exemplar condition for same-direction trials (*t* (18)= 2.77, *p*_FDR_ = 0.04). A two-way ANOVA revealed a significant main effect of motion direction (*F* (1,18)= 13.10, *p* = 0.002, *η*^2^ = 0.42), but no significant main effect of category (*F* (3,54)= 1.17, *p* = 0.33, *η*^2^ = 0.06), or interaction between category and direction (*F* (3,54)= 0.29, *p* = 0.83, *η*^2^ = 0.02). These results suggest that surround suppression in hMT+ depends primarily on the relative motion direction between center and surround.

**Figure 2:**
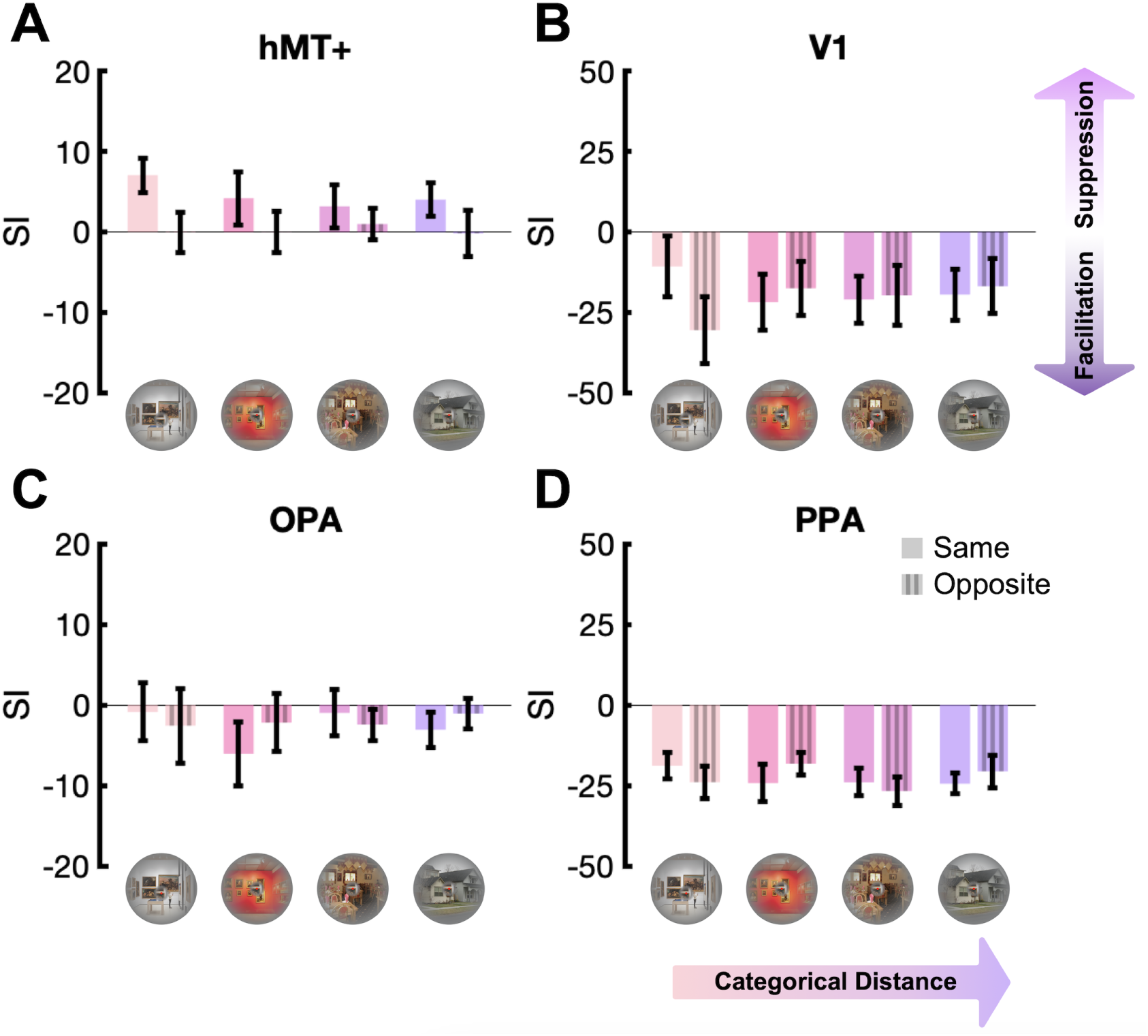
Univariate surround modulation effects. Suppression index (SI) values are shown across four category conditions: identical exemplar, different exemplar, different basic-level category, and different superordinate category, and two motion-direction conditions (same and opposite). Example stimuli for each category condition are displayed along the x-axis to illustrate the categorical hierarchy, with categorical distance increasing from left to right (from the identical exemplar to the different superordinate category conditions). Results are displayed for **A)** hMT+, **B)** V1, **C)** OPA, and **D)** PPA. Positive SI values indicate surround suppression; negative values reflect facilitation. Error bars represent the standard error of the mean (SEM).

As expected, the categorical similarity of the center and surround had no effect on the suppression strength, consistent with hMT+ being primarily motion selective rather than sensitive to scene content. Because the categorical similarity neither modulated SI nor interacted with motion direction, SI values were averaged across the four category conditions (Figure 3A). A paired-sample *t*-test on the averaged data confirmed that suppression was significantly stronger when the center and surround moved in the same direction than when they moved in opposite directions (*t*(18) = 3.62, *p* = 0.002). Together, these results demonstrate that hMT+ exhibits motion-dependent surround suppression under dynamic, natural scene conditions, consistent with perceptual suppression effects in our recent behavioral study using the same stimuli (Kiniklioglu & Kaiser, 2025), and with classical findings using simple gratings (Kiniklioglu & Boyaci, 2022; Paffen, Alais, & Verstraten, 2005; Paffen, van der Smagt, et al., 2005).

**Figure 3:**
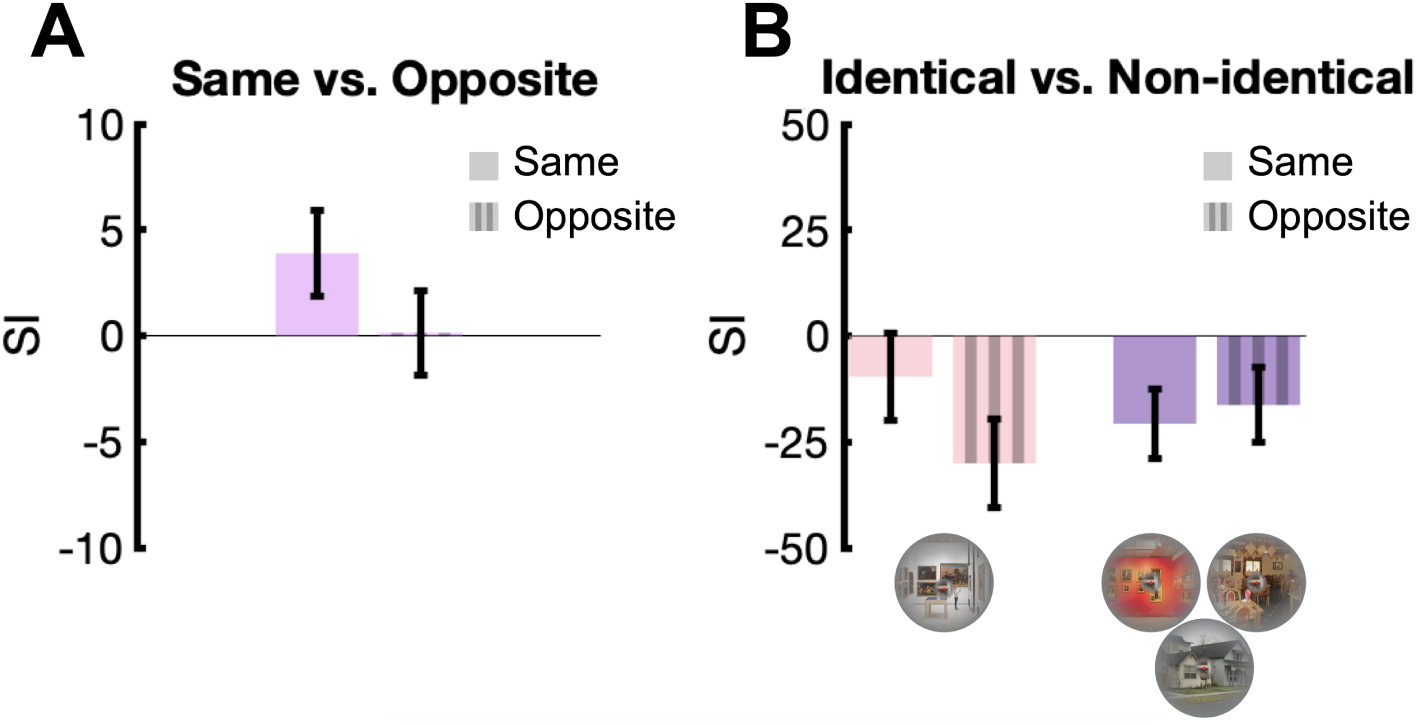
Averaged univariate surround modulation effects in hMT+ and V1. **A)** hMT+ SI values averaged across the four category conditions for sameand oppositedirection trials. **B)** V1 SI values for the identical exemplar condition versus the mean of the three other (non-identical) conditions (i.e., different exemplar, different basic-level category, and different superordinate category conditions), shown separately for same- and opposite-direction trials. Positive SI values indicate surround suppression; negative values reflect facilitation. Error bars represent the standard error of the mean (SEM).

In V1, we observed surround facilitation across all conditions, in stark contrast to the suppression effects found in hMT+. Figure 2B shows the SI values for the V1 sub-ROI across four scene-category conditions and two motion-direction conditions. SI values were significantly lower than zero in all conditions except the identical-exemplar condition for same-direction trials (*t* (18)= 0.95, *p*_FDR_ = 0.18). A two-way ANOVA revealed a significant interaction between motion direction and categorical similarity (*F* (3,54)= 9.34, *p <* 0.001, *η*^2^ = 0.34), but no significant main effects of motion direction (*F* (1,18)= 0.44, *p* = 0.51, *η*^2^ = 0.02) or category (*F* (3,54)= 0.30, *p* = 0.83, *η*^2^ = 0.02).

Because the three non-identical category conditions (different exemplar, different basic-level category, and different superordinate category) did not differ from each other and showed no effect of motion direction (all *p*s*>*0.20), they were collapsed into a single “non-identical” condition, analyzed separately for same- and opposite-direction trials (Figure 3B). Post-hoc tests confirmed that the category × motion direction interaction observed in the ANOVA was driven by the identical-exemplar condition. Motion direction significantly affected responses only in the identical-exemplar condition, with reduced facilitation when the center and surround moved in the same direction compared to opposite directions (*t*(18) = 3.47, *p*_FDR_ = 0.003). Moreover, for same-direction trials, facilitation was significantly weaker for identical exemplars than for the collapsed non-identical condition (*t*(18) = 2.78, *p*_FDR_ = 0.01). This pattern suggests that similarity between center and surround modulates facilitation in V1, such that identical center–surround combinations yield weaker facilitation than non-identical ones, even within the same category. This result aligns with classical low-level findings showing the strongest suppression for iso-oriented uniform stimuli (Cavanaugh et al., 2002a; DeAngelis et al., 1994; Flevaris & Murray, 2015; Schallmo et al., 2016; Serrano-Pedraza et al., 2012; Walker et al., 1999) and is consistent with motion-dependent surround modulation observed in hMT+, though in V1 the effect manifested as reduced facilitation rather than suppression.

Scene-selective areas OPA and PPA both showed surround facilitation. Figures 2C and D show the SI values for these ROIs across four scene-category conditions and two motion-direction conditions. In OPA, SI values trended toward facilitation but did not reach significance (all *p*_FDR_ *>* 0.05), except the different-exemplar (*t* (18)= 2.44, *p*_FDR_ = 0.03) and different superordinate category condition (*t* (18)= 2.52, *p*_FDR_ = 0.03) for same-direction trials. In contrast, PPA showed robust facilitation, with SI values significantly below zero across all conditions (all *t* (18)*>* 3.76, *p*_FDR_ *<* 0.001). A two-way ANOVA in both OPA and PPA revealed no significant effects of motion direction, category, or their interaction (all *p*s *>* 0.05). These results contrast with the suppression observed in hMT+ and the reduced facilitation in V1 under identical-exemplar condition, suggesting that scene-selective regions rely on distinct surround modulation mechanisms under dynamic naturalistic stimulation. These higher-level scene areas may integrate information from both center and surround to support scene understanding rather than exhibiting classical surround suppression.

### Multivariate analyses reveal motion-dependent surround modulation in hMT**+**

To assess how motion direction influences surround modulation, data were collapsed across the four category conditions prior to decoding to isolate motion-related effects. We then decoded the center-only versus center+surround conditions in V1, hMT+, OPA, and PPA under two motion-direction conditions: same-direction and opposite-direction.

As shown in Figure 4A, decoding accuracy in hMT+ was above chance for both same-direction (*p*_FDR_ *<* 0.001) and opposite-direction trials (*p*_FDR_ = 0.003). A paired-samples *t*-test revealed higher accuracy for same-direction than opposite-direction trials (*t*(18) = 3.07, *p* = 0.007), indicating that motion congruence enhances surround modulation in hMT+ multivoxel responses. This multivariate pattern closely mirrors the univariate results, which showed stronger surround suppression when center and surround moved in the same direction. Together, these analyses demonstrate that hMT+ activity reflects motion-dependent surround modulation both in overall response amplitude and in distributed activation patterns. These converging findings suggest that surround modulation in hMT+ is driven primarily by the relative motion direction between center and surround, consistent with classical low-level results (Kiniklioglu & Boyaci, 2022; Paffen, Alais, & Verstraten, 2005; Paffen, van der Smagt, et al., 2005).

**Figure 4:**
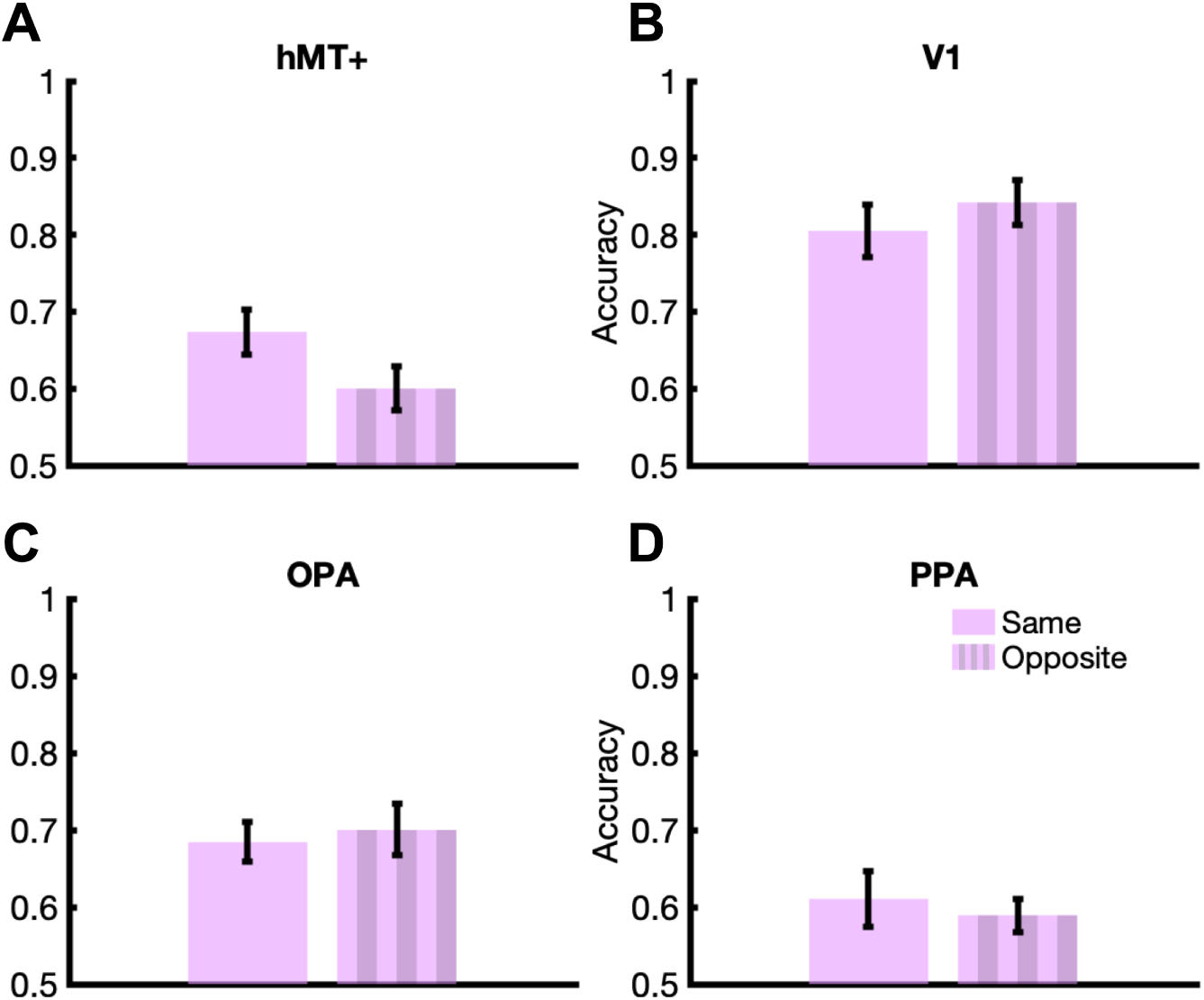
Multivoxel decoding accuracy for center-only and center+surround classification for the two motion-direction conditions: same-direction and opposite-direction. Mean decoding accuracy is shown for **A)** hMT+, **B)** V1, **C)** OPA, and **D)** PPA. Error bars indicate the standard error of the mean (SEM).

As shown in Figure 4B–D, decoding accuracy was above chance in V1, OPA, and PPA for both same-direction (all *p*_FDR_ *<* 0.008) and opposite-direction trials (all *p*_FDR_ *<* 0.001). However, paired-samples *t*-tests revealed no significant differences in decoding accuracy between same- and opposite-direction trials (all *p*_FDR_ *>* 0.20). Although V1, OPA, and PPA showed reliable above-chance decoding, the absence of direction-related differences in these regions suggests that their surround modulation reflects general contextual integration rather than motion congruence.

### Multivariate analyses reveal category-dependent surround modulation in OPA and PPA

To assess how category similarity influences surround modulation, data were collapsed across the two motion-direction conditions prior to decoding. We then decoded the center-only versus center+surround conditions in V1, hMT+, OPA, and PPA across four category conditions: identical exemplar, different exemplar, different basic-level category, and different superordinate category.

As shown in Figures 5A and B, classification accuracy was significantly above chance for all category conditions in both hMT+ (all *p*_FDR_ *<* 0.01) and V1 (all *p*_FDR_ *<* 0.001). Despite robust decoding performance, one-way repeated-measures ANOVAs revealed no effect of category in either region (V1: *F* (3, 54) = 1.93, *p* = 0.13, *η*^2^ = 0.10; hMT+: *F* (3, 54) = 0.56, *p* = 0.64, *η*^2^ = 0.03). These results indicate strong surround modulation in both areas, but this modulation did not depend on categorical similarity between center and surround, consistent with the roles of V1 and hMT+ in processing low-level visual features such as edges, orientation, contrast, and motion rather than higher-level, category-selective information.

**Figure 5:**
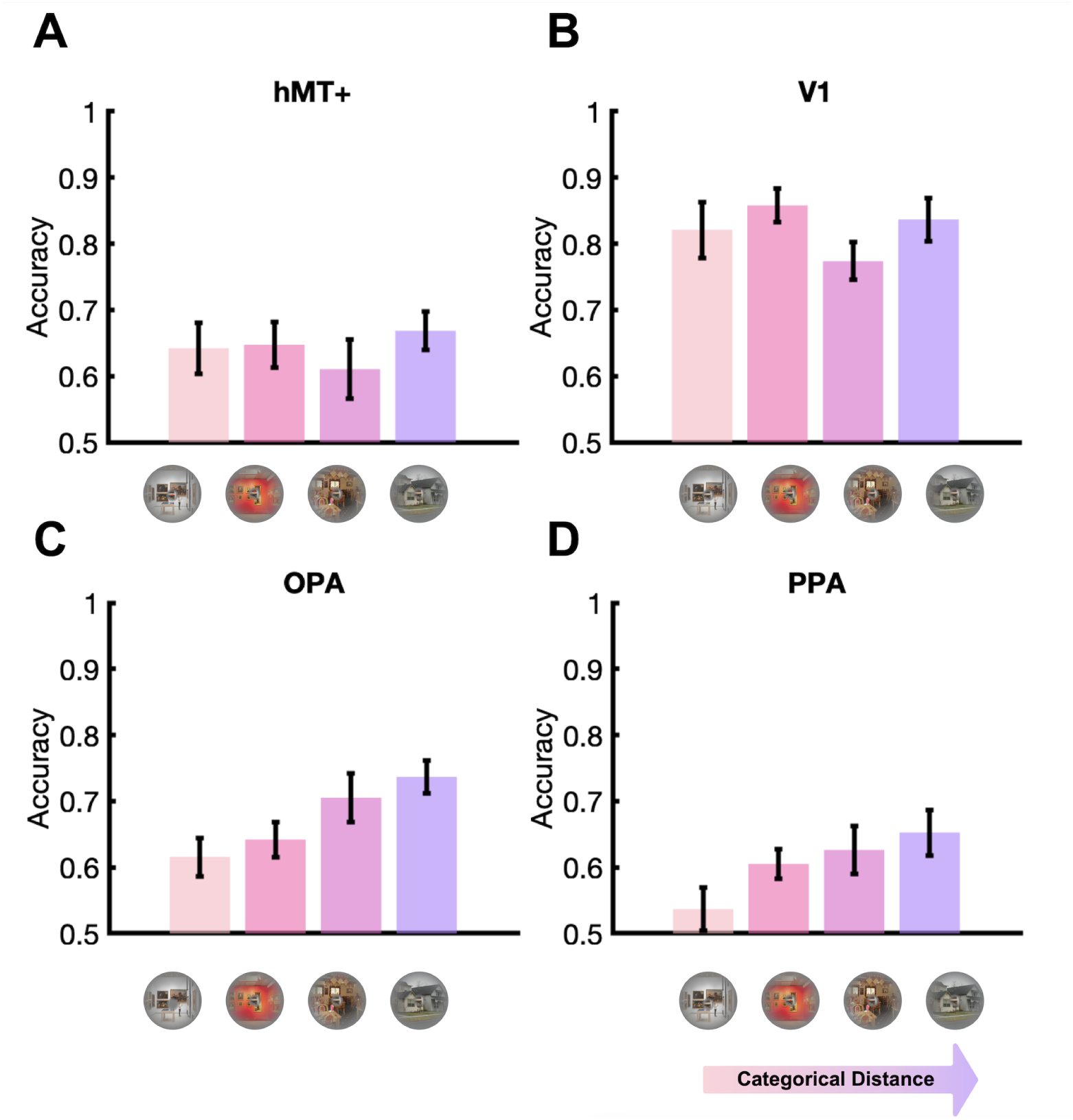
Multivoxel decoding accuracy for the center-only versus center+surround classification across the four category conditions: identical exemplar, different exemplar, different basic-level category, and different superordinate category. Example stimuli for each category condition are displayed along the x-axis to illustrate the categorical hierarchy, with categorical distance increasing from left to right (from the identical exemplar to the different superordinate category conditions). Mean decoding accuracy is shown for **A)** hMT+, **B)** V1, **C)** OPA, and **D)** PPA. Error bars represent the standard error of the mean (SEM).

As shown in Figure 5C, the classification accuracy was significantly above chance in OPA for all category conditions (all *p*_FDR_ *<* 0.002). A one-way repeated-measures ANOVA revealed a main effect of category, *F* (3, 54) = 4.12, *p* = 0.01, *η*^2^ = 0.19. Posthoc comparisons showed higher decoding for the different superordinate-category condition than for both the identical- and different-exemplar conditions (all *t*(18) *>* 3.01, *p*_FDR_ *<* 0.02), and marginally higher decoding for the different basic-level category condition than for the identical-exemplar condition (*t*(18) = 2.04, *p*_FDR_ = 0.06). Decoding did not differ between the identical- and different-exemplar conditions, *t*(18) = 0.70, *p*_FDR_ = 0.51.

These results suggest that surround modulation in OPA increases with categorical distance. Multivoxel responses were more strongly influenced by the surround when the center and surround scenes belonged to different categories, indicating that OPA is sensitive not only to the presence of contextual surround input but also to its semantic relationship with the center. This categorical sensitivity aligns with our behavioral findings (Kiniklioglu & Kaiser, 2025), in which surround suppression increased as categorical similarity decreased.

PPA showed a similar pattern, with decoding accuracy increasing as the categorical dissimilarity between center and surround increased. Decoding was above chance for all category conditions except the identical-exemplar condition (*p*_FDR_ = 0.17; Figure 4D).

The main effect of category was significant, *F* (3, 54) = 2.93, *p* = 0.04, *η*^2^ = 0.14. Posthoc comparisons revealed higher decoding accuracy for the different superordinate-category condition than for the identical-exemplar condition (*t*(18) = 2.59, *p* = 0.02), and higher decoding accuracy for the different basic-level category condition than for the identical-exemplar condition (*t*(18) = 2.18, *p* = 0.04). Compared with OPA, surround modulation in PPA exhibited a more gradual and weaker increase in decoding accuracy with categorical distance. Taken together, these results extend surround modulation effects to the scene-selective cortex, demonstrating that surround modulation in these regions is shaped by higher-level categorical relationships.

## Discussion

The present study investigated how motion congruence and categorical similarity between center and surround scenes shape neural surround modulation. Using fMRI, we measured neural responses while systematically varying the relationship of center and surround across four levels of categorical similarity (identical exemplar, different exemplar, different basic-level category, and different superordinate category) and two levels of motion congruence (same vs. opposite direction). Univariate analyses revealed distinct patterns of surround modulation across the visual hierarchy. hMT+ showed robust motion-dependent surround suppression, with stronger suppression when the center and surround moved in the same direction. In contrast, V1 exhibited surround facilitation across all conditions, which was reduced when the center and surround were identical and moved in the same direction. Scene-selective regions OPA and PPA also showed predominantly facilitation rather than suppression. Multivariate analyses complemented these findings: in hMT+, decoding between the center-only versus center+surround conditions was stronger for samethan opposite-direction motion, whereas in OPA and PPA, surround modulation was category-dependent, with decoding accuracy increasing as categorical similarity between center and surround decreased.

These results provide the first neural evidence for surround modulation in humans with dynamic natural scenes. While surround suppression has been extensively characterized with simple stimuli such as gratings (Cavanaugh et al., 2002b; Schallmo et al., 2018; Tadin et al., 2003), its neural basis under naturalistic conditions has remained largely unexplored. The present findings demonstrate that surrounding scene context systematically modulates neural responses to central scenes across multiple stages of the visual hierarchy: from low-level facilitation in V1 to motion-dependent suppression in hMT+ and category-dependent modulation in scene-selective regions. This hierarchical organization suggests that surround modulation, a well-established mechanism in early visual processing, is also engaged during naturalistic scene perception, with both motion congruence and categorical similarity shaping how contextual information is integrated across the visual system.

### Motion-dependent surround modulation in hMT**+**

hMT+ exhibited stronger suppression when the center and surround moved in the same direction than when they moved in opposite directions. Decoding analyses further confirmed that this surround modulation was driven specifically by the motion direction, independent of categorical similarity. This pattern aligns with classical neurophysiological findings showing that MT neurons show stronger suppression for same-direction motion and weaker or no suppression when the center and surround move in opposite directions (Allman et al., 1985; Born & Tootell, 1992; Cavanaugh et al., 2002b; Kastner et al., 1995; Lamme, 1995). Such motion-dependent modulation likely reflects local inhibitory interactions among direction-selective neurons in MT, consistent with the *MT hypothesis* of surround suppression (Tadin et al., 2003). By extending these low-level findings to naturalistic conditions, our results provide neural evidence that motion-based center–surround interactions are recruited during complex scene processing in hMT+. Furthermore, the close correspondence between our neural and behavioral findings using identical stimuli (Kiniklioglu & Kaiser, 2025) suggests that perceptual suppression during natural vision may arise from these motion-sensitive inhibitory mechanisms in hMT+.

### Facilitation in V1 Reflects Sensitivity to Physical Similarity

Although facilitation in V1 was robust across conditions, it was significantly weaker for identical exemplars than for other category conditions in same-direction trials, indicating that greater physical similarity between the center and surround reduces facilitation. Moreover, motion direction affected responses only in the identical-exemplar condition, with facilitation decreasing when the center and surround moved in the same direction compared to opposite directions. This pattern aligns with classical findings showing the strongest suppression for iso-oriented, uniform stimuli (Cavanaugh et al., 2002a; DeAngelis et al., 1994; Flevaris & Murray, 2015; Schallmo et al., 2016; Serrano-Pedraza et al., 2012; Walker et al., 1999). The motion-dependent modulation also resembles suppression effects reported in hMT+; however, in V1, the influence manifests as reduced facilitation rather than suppression. These effects may arise from horizontal interactions within V1 or feedback from motion-selective areas such as hMT+, consistent with evidence that motion information can be transmitted across early visual areas (Angelucci & Bressloff, 2006; Angelucci et al., 2017; Hupé et al., 2001).

Supporting this interpretation, animal studies using naturalistic stimuli have shown that surround modulation in the primary visual cortex depends strongly on low-level image similarity. For example, Coen-Cagli et al. (2015) reported that macaque V1 exhibits stronger suppression when the center and surround consist of homogeneous natural image regions. This aligns broadly with our findings, which also reveal surround modulation driven by physical similarity, although it manifested as reduced facilitation rather than increased suppression. Consistent with this, multivariate analyses further showed robust surround modulation in V1, but this effect did not depend on categorical similarity or motion congruence. Together, these results indicate that surround modulation in V1 primarily reflects sensitivity to low-level physical similarity between center and surround, rather than higher-level, category-selective information. Nevertheless, the nature of surround modulation in V1 remains debated. Some studies have reported suppression, suggesting that early visual mechanisms may contribute to the suppressive effects often observed in higher-level visual areas (Angelucci et al., 2017; Nurminen et al., 2009, 2013; Zenger-Landolt & Heeger, 2003), whereas others have reported surround facilitation (Er et al., 2020; Press et al., 2001). Importantly, several studies have demonstrated both suppression and facilitation in V1, depending on stimulus configuration, task demands, and attentional context (Flevaris & Murray, 2015; Ichida et al., 2007; Williams et al., 2003). It is therefore possible that the facilitation observed in our study reflects context-dependent modulation. Because the task did not explicitly require attention to the center–surround relationship, suppressive interactions may have been reduced, resulting in an overall facilitative response.

Alternatively, the facilitation observed in V1 may be influenced by feedback from scene-selective cortex. Surround modulation in V1 is thought to arise from a complex interplay of feedforward, horizontal, and feedback mechanisms (Nurminen & Angelucci, 2014; Nurminen et al., 2018). As part of this highly interconnected network, V1 receives extensive feedback from other visual areas (Shao & Burkhalter, 1996), including those associated with scene processing (Morgan et al., 2019; Rockland & Van Hoesen, 1994). Top-down inputs from scene-selective regions may bias early visual responses toward facilitation, dynamically regulating the balance between facilitation and suppression and thereby contributing to the overall pattern observed in V1.

### Category-dependent modulation in scene-selective cortex

OPA and PPA primarily exhibited facilitation, likely reflecting their relatively large receptive fields that integrate information across broad regions of the visual field (e.g., Levy et al., 2001; Nasr et al., 2011; Silson et al., 2015). This integration may allow them to combine inputs from both center and surround to support scene understanding rather than exhibiting classical surround suppression. While univariate analyses revealed no category or motion effects, the more sensitive multivariate analyses showed clear category-dependent surround modulation, particularly in OPA. Multivoxel responses were more strongly influenced by the surround when the center and surround scenes belonged to different categories, indicating that OPA is sensitive not only to the presence of contextual information but also to its categorical relationship with the central scene. PPA showed a similar but weaker gradual increase in surround modulation with categorical distance, possibly reflecting the lower signal-to-noise ratio typically observed in ventral visual regions (e.g., Rua et al., 2018; Winawer et al., 2010).

Importantly, decoding revealed no motion-related effects in either region, indicating that surround modulation in scene-selective cortex is driven specifically by categorical similarity. These results are consistent with our behavioral findings (Kiniklioglu & Kaiser, 2025), suggesting a shared organizing principle across neural and perceptual levels whereby contextual influences strengthen as scenes become more categorically dissimilar. The correspondence between neural and behavioral results obtained with identical stimuli supports the idea that perceptual contextual effects during natural scene perception may reflect category-sensitive integrative processes in scene-selective regions.

Similar sensitivity to categorical context has been demonstrated in prior neuroimaging work on scene and object perception. Contextual incongruence between scene elements has been shown to alter neural responses in parahippocampal and occipitotemporal regions (Faivre et al., 2019; Peyrin et al., 2021; Ŕemy et al., 2014). For example, Peyrin et al. (2021) reported that categorically congruent (e.g., both man-made) but physically dissimilar (e.g., buildings vs. streets) peripheral scenes disrupted central scene categorization and increased activation in inferior frontal and occipitotemporal cortices. Consistent with this, we found that surround modulation in scene-selective cortex was stronger even when the center and surround belonged to the same superordinate category (e.g., two indoor scenes) but differed at the basic level (e.g., restaurant vs. museum).

### Hierarchical surround modulation may support efficient scene representation

Real-world scenes show systematic spatial and semantic regularities (e.g., a bathroom typically contains a sink, mirror, and shower arranged within a characteristic layout). The visual system exploits these regularities to integrate information efficiently across multiple representational levels, supporting rapid understanding of complex environments (Bar, 2004; Kaiser et al., 2019; Võ, 2021). Computational and neurophysiological models suggest that such contextual integration relies on recurrent and inhibitory interactions that predictively modulate sensory input according to its relevance (Angelucci & Bressloff, 2006; Gilbert & Li, 2013; Rao & Ballard, 1999). Consistent with this view, behavioral and neuroimaging work demonstrates that coherent scene structure facilitates both scene and object perception (Chen et al., 2022; Davenport & Potter, 2004; Kaiser & Peelen, 2018; Kaiser et al., 2019, 2020, 2021; Võ & Wolfe, 2013). Together, these findings suggest that contextual mechanisms may enhance informative signals while attenuating redundant input, thereby promoting stable and efficient visual processing in natural environments.

Our results build on this framework by showing that categorically incongruent surrounds elicit stronger contextual modulation in scene-selective cortex than congruent surrounds. Whereas studies using simple stimuli have attributed surround modulation to the suppression of statistical redundancies in low-level features (Angelucci et al., 2017; Coen-Cagli et al., 2012; Nurminen & Angelucci, 2014; Schwartz & Simoncelli, 2001; Vinje & Gallant, 2000), our findings indicate that, at higher representational levels, contextual influences are governed primarily by categorical relationships. In natural scenes, categorically incongruent information may enhance neural responses because it violates contextual expectations and conveys higher informational value, whereas with simple stimuli, identical information may be suppressed because it is predictable and redundant. In contrast, V1 showed a pattern more consistent with low-level surround modulation, where facilitation decreased when the center and surround were physically identical, indicating sensitivity primarily to visual redundancy rather than categorical context. Both forms of modulation may therefore reflect a shared computational principle: optimizing neural representations by emphasizing informative differences while attenuating predictable input.

Together, these results reveal a hierarchical organization of surround modulation in natural vision. Early visual areas (V1) are primarily governed by physical similarity, mid-level motion areas (hMT+) are sensitive to motion congruence, and higher-level scene-selective regions (OPA and PPA) integrate categorical structure. This progression supports a predictive and efficient coding framework (Angelucci et al., 2017; Gilbert & Li, 2013; Rao & Ballard, 1999), whereby each stage of the visual hierarchy selectively suppresses predictable or redundant input and enhances informative discrepancies.

## Limitations and future directions

Although the present study provides neural evidence for both categorical and motion-based surround modulation in natural vision, several questions remain open for future investigation. First, we did not explicitly manipulate low-level visual features such as spatial frequency or amplitude spectrum, which may covary with categorical similarity and contribute to surround modulation (Peyrin et al., 2021). Future work could examine these factors more directly by varying physical properties and categorical coherence independently to disentangle their respective influences.

Second, while we used sub-ROI localizers in V1 and hMT+ to identify voxels corresponding to the center stimulus, this approach could not be applied to OPA and PPA due to their larger receptive fields. Consequently, the ROIs in these regions may have included voxels responsive to both center and surround, so that the effects cannot be directly interpreted as modulatory effects on the representation of the central stimulus. Moreover, scene-selective areas are thought to contain fine-grained functional subfields with heterogeneous tuning to spatial scale, visual field position, and category information (e.g., Baldassano et al., 2013; Silson et al., 2015). The absence of univariate category effects in OPA and PPA may therefore reflect this spatial and functional heterogeneity, where the averaging across voxels with different category or positional preferences could mask category-specific responses. Category-specific modulatory effects may instead depend on multivariate response patterns in scene-selective cortex, necessitating pattern-based analysis approaches for characterizing this fine-scale organization.

## Conclusions

Using dynamic natural scenes, we demonstrate that surround modulation is a canonical property of human visual processing, shaped by both motion-direction congruence and categorical similarity. Univariate and multivariate analyses revealed distinct mechanisms across the visual hierarchy: in hMT+, surround suppression depended on motion congruence (stronger for same-direction motion), consistent with classic center–surround interactions among direction-selective neurons; in V1, responses were predominantly facilitatory but weakened when center and surround were physically identical and moved in the same direction, reflecting sensitivity to low-level similarity; in scene-selective cortex (OPA, PPA), surround influences were largely facilitatory and scaled with categorical distance in multivoxel patterns, indicating sensitivity to higher-level categorical structure. Together, these findings extend surround modulation from simple stimuli to naturalistic vision, with contextual influences shifting from sensitivity to physical similarity in the early visual cortex to category-sensitive integration in higher-level regions. This hierarchical progression aligns with predictive and efficient coding accounts, whereby each stage attenuates predictable input and emphasizes informative discrepancies to support perceptual stability and efficient scene understanding.

## Data Availability

The data set and analysis scripts will be available on the Open Science Framework.

## Acknowledgements

This work was supported by the Deutsche Forschungsgemeinschaft (DFG), grants SFB/TRR 135 (project no. 222641018); KA4683/5-1 (project no. 518483074); and under Germany’s Excellence Strategy (EXC 3066/1, “The Adaptive Mind”, project no. 533717223). It was further supported by an European Research Council (ERC) Starting Grant (PEP, ERC-2022-STG 101076057). Views and opinions expressed are those of the authors only and do not necessarily reflect those of the European Union or the European Research Council. Neither the European Union nor the granting authority can be held responsible for them. MR imaging for this study was performed at the Bender Institute of Neuroimaging (BION) at Justus Liebig University Giessen, Germany. We thank Tugce Dalmis for assistance with data collection.

